# *In silico* characterization of mechanisms positioning costimulatory and checkpoint complexes in immune synapses

**DOI:** 10.1101/2020.01.16.908723

**Authors:** Anastasios Siokis, Philippe A. Robert, Philippos Demetriou, Audun Kvalvaag, Salvatore Valvo, Viveka Mayya, Michael L. Dustin, Michael Meyer-Hermann

**Affiliations:** Department of Systems Immunology and Braunschweig Integrated Centre of Systems Biology, Helmholtz Centre for Infection Research, Braunschweig, 38106, Germany; Institute of Biochemistry, Biotechnology and Bioinformatics, Technische Universität Braunschweig, Braunschweig, 38106, Germany; Kennedy Institute of Rheumatology, Nuffield Department of Orthopaedics, Rheumatology and Musculoskeletal Sciences, University of Oxford, Oxford OX3 7FY, United Kingdom; Department of Molecular Cell Biology, Institute for Cancer Research, Oslo University Hospital, Montebello, N-0379 Oslo, Norway

**Author notes:** Department of Immunology, University of Oslo, Oslo, 0372 Norway (Philippe A. Robert).

## Abstract

Integrin and small immunoglobulin superfamily (sIGSF) adhesion complexes function physiologically in human immunological synapses (IS) wherein sIGSF complexes form a corolla of microdomains around an integrin ring and secretory core. The corolla recruits and retains the major costimulatory and checkpoint complexes that regulate the response to T cell receptor (TCR) engagement, making forces that govern corolla formation of particular interest. We developed a phenomenological agent-based model in order to test different hypotheses concerning the mechanisms underlying molecular reorganization during IS formation. The model showed that sIGSF complexes are passively excluded to the distal aspect of the IS as long as their interaction with the ramified F-actin transport network is absent or weaker than that of integrins. An attractive force between sIGSF adhesion and costimulatory/checkpoint complexes relocates the latter from the centre of the IS to the corolla. The simulations suggest that size based sorting interactions with large glycocalyx components as well as a short-range self-attraction between sIGSF complexes explain the corolla “petals”. These molecular and mechanistic features establish a general model that can recapitulate complex pattern formation processes observed in cell-bilayer and cell-cell interfaces.

**One Sentence Summary:** Computer simulations of immunological synapses reveal the localization mechanisms of immunoglobulin superfamily adhesion and costimulatory/checkpoint complexes.

## Introduction

Immunological synapse (IS) organization is important for signal integration and effector function of T cells (*1-4*). In humans, the immune synapse is built around T cell receptor (TCR) interaction with peptide-major histocompatibility complex (pMHC) to form a specificity complex and integrin and small immunoglobulin superfamily (sIGSF) adhesion complexes. The integrin LFA-1 (lymphocyte function-associated antigen 1) form complexes with intercellular adhesion molecule-1 (ICAM-1) and sIGSF member cluster of differentiation 2 (CD2) with human CD58 (hCD58) in the context of T cell effector function (*5*). Integrin-mediated adhesion is an active process regulated by interactions with the F-actin transport network (*6*), whereas the sIGSF members operate through highly multivalent microdomains and are less dependent on cellular energy (*7*). When a live T cell forms an interface with a planar supported lipid bilayer (SLB) containing freely mobile ICAM-1, CD58 and pMHC, the individual complexes are organized into a layered, radially symmetrical pattern with central TCR-pMHC cluster (synaptic cleft and central supramolecular activation cluster, cSMAC), surrounded by an integrin adhesion ring (peripheral or pSMAC), and fringed by sIGSF microdomains in the distal SMAC (dSMAC), resembling flower petals known as corolla (5, 8*-11*). cSMAC and pSMAC formation driven by F-actin mediated centripetal transport (*10, 12, 13*) and intermittent coupling of TCR-pMHC to the F-actin network (*14*) with a constant force (*15*). The corolla pattern is completely distinct from that formed by costimulatory and checkpoint complexes, like CD28-CD80 and PD1-PDL1, which also occupy TCR microclusters and the outer annulus of the cSMAC (*3, 11, 16-18*). We have previously shown that cSMAC localization of both TCR-pMHC and costimulatory complexes can be simulated when engaged TCR and CD28 interact independently with the F-actin transport network (*19*). Strikingly, addition of CD2-CD58 complexes to this system leads to corolla formation and relocation of the costimulatory complexes to the corolla in the dSMAC (*11*). Thus, it is important to understand the forces that regulate corolla formation.

Interactions in the IS display phase separation behavior based on chemical kinetics coupled to membrane spacing (*20*). CD2 interaction with mouse CD48 (mCD48) assembles adhesion junctions with an intermembrane gap of 12.8 nm (*7, 21*), similar to TCR-pMHC interaction at 13.1 nm (*22*). CD2 interactions with hCD58 or mCD48 are low affinity with fast dissociation kinetics (*23-26*). The long, unstructured and highly conserved cytoplasmic domain of CD2 is rich in polyproline motifs (PP) that contribute to Src family kinase activation (*10, 11, 27-30*). Similar unstructured domains with multiple PP in combination with proteins containing multiple SH3 domains can undergo concentration dependent liquid-liquid phase separation (*31*). CD2-CD58 complex formation supports T cell polarization (*32*), and also augments and sustains antigen-induced cytoplasmic calcium (Ca^2+^) increase in the T cell (*33*). Blockade of the CD2-CD58 interaction has been shown to impair recruitment of PLCγ1, a key player for down-stream signaling in T cells (*33*).

Together with the above-mentioned molecules, T cells and antigen presenting cells (APCs) express numerous other surface glycoproteins that make up the glycocalyx, which plays important roles in T cell interactions, activation and effector function. A characteristic glycocalyx protein is the transmembrane protein tyrosine phosphatase CD45. No ligand has been described for CD45 on APCs. Nevertheless, it plays an important role for TCR signaling (*34*), positively regulating T cell activation via dephosphorylation of the inhibitory C-terminal tyrosine of p56lck and p59fyn proteins (*35-38*), which also interact with CD2 (*27*). Importantly, CD45 molecules are excluded from the TCR-pMHC microclusters during the initial moments of IS formation and eventually completely relocate to the outer region of the IS, the dSMAC (*35, 39, 40*), where the corolla forms (*11*). This suggests that CD45 may play a role in the organization of the corolla/dSMAC, although no previous studies have addressed this question to the best of our knowledge.

In this study, we focused on understanding the possible mechanisms leading to the adhesion corolla in the dSMAC. We developed a general agent-based model platform for the simulation of immunological synapse formation including all relevant components like specificity, intergrin, sIGSF adhesion and costimulatory/checkpoint complexes as well as glycocalyx molecules. This complete platform reflects physical processes like diffusion, chemical kinetics and agent-agent interactions. Size-based segregation (SBS) was modeled as a repulsive force between differently sized complexes and F-actin centripetal transport as an empirical centrally directed force (*19*). In the simulations, an LFA-1 gradient develops in the pSMAC. This implies that the CD2-CD58 complexes comprise two distinct populations, one passively following the TCR-pMHC movement towards the cSMAC and the other relocated to the dSMAC and residing in an annular pattern. Attraction between sIGSF adhesion and costimulation/checkpoint complexes resulted in the formation of the corolla pattern, where the two classes of interactions can work together. In order to recapitulate the corolla in the absence of costimulatory interactions, a self-attraction between sIGSF adhesion complexes was sufficient. The origin of such a force could be the interaction of sIGSF adhesion complexes with local F-actin asters and focal points but not the centripetal F-actin flow. In simulations with a glycocalyx component present, the corolla pattern was reproducible alone by the SBS between sIGSF adhesion complexes and glycocalyx molecules. Taken together, these results imply that the corolla pattern formation can be a result of an active process, such as the sIGSF self-attraction, or a physical phase separation process together with competition for space in the presence of glycocalyx components.

## Results

### sIGSF adhesion complexes alter CD28 localization

In our effort to understand the mechanisms underlying sIGSF adhesion complex localization and interaction with other molecules on the forming synapse, such as costimulatory/checkpoint complexes, we developed a phenomenological agent-based model and validated it by experiments performed on SLBs (*11, 16*). This model takes into account chemical kinetics for the formation of receptor-ligand complexes. SBS of sIGSF versus longer integrin complexes is modeled as a repulsive force and the F-actin driven centripetal flow of complexes as a centrally directed force (Fig. 1). As those interactions depend on the biophysical properties of the complexes like size and on- and off-rates, the following results were generated for a specific example of the involved complexes: The specificity complex TCR-pMHC, the integrin complex LFA-1-ICAM-1, costimulation/checkpoint complexes CD28-CD80 (similar to PD1-PDL1), and sIGSF adhesion complex CD2-CD58.

**Fig. 1.**
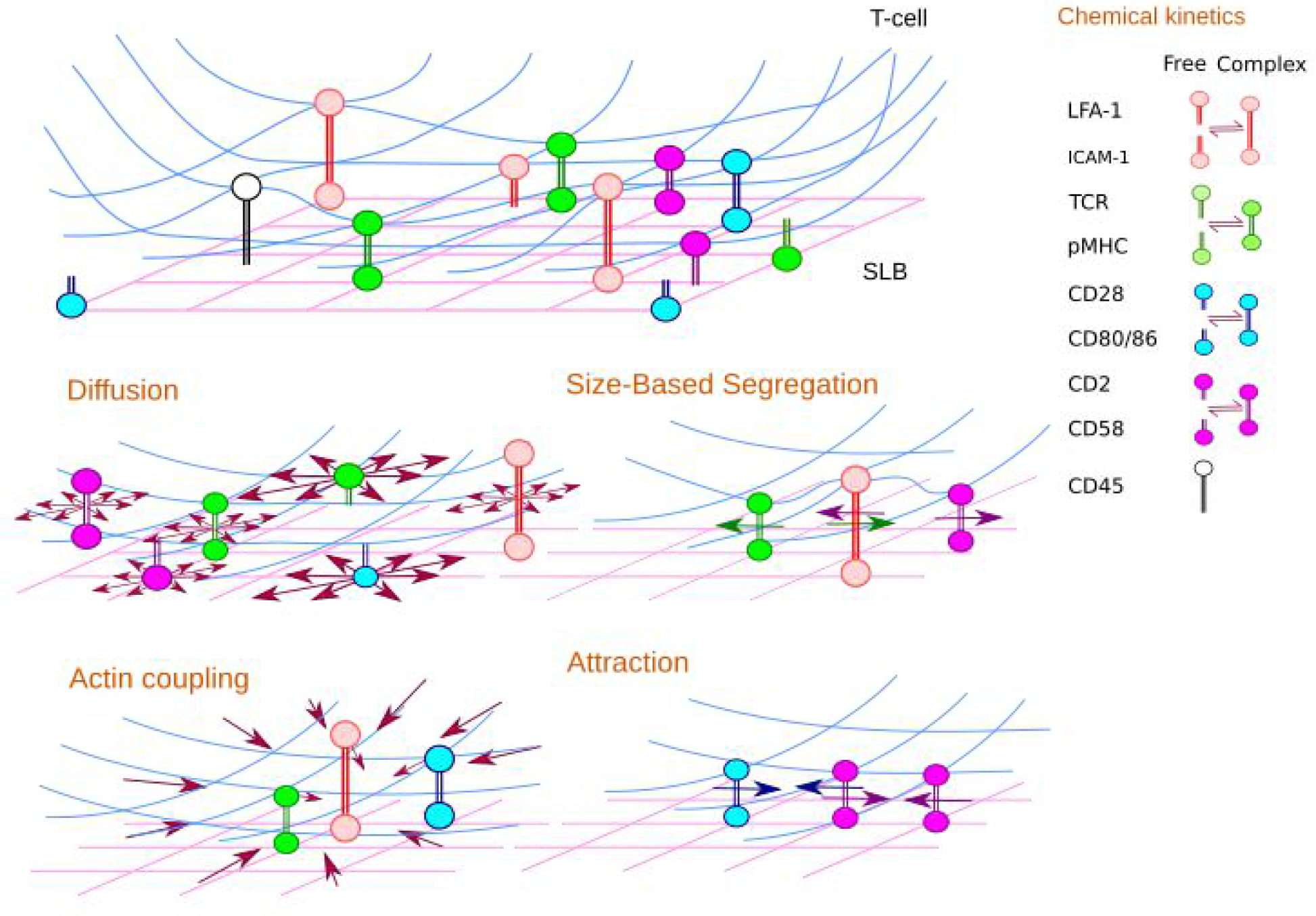
List of mechanisms included in the IS model. The membrane of the T cell (blue) and the SLB (pink) carry molecules. Opposite molecules bind or unbind by chemical kinetics. Free and complexed molecules move by diffusion or forces: centripetal forces reflecting F-actin coupling, SBS representing the effect of membrane bending and attractive forces representing interaction with local F-actin asters or of other origin.

SBS between TCR-pMHC and LFA-1-ICAM-1 complexes was responsible for the formation of TCR-pMHC microclusters, while at the same time the F-actin driven centripetal force results in the accumulation of TCR-pMHC in the cSMAC as well as the emergence of an LFA-1-ICAM-1 gradient in the pSMAC (Fig. 2A-top row). The gradient acted as an exclusion mechanism, altering the localization of free molecules as well as receptor-ligand complexes, which did not interact with the F-actin centripetal flow. Costimulatory CD28-CD80 complexes (*16*) are structurally similar to the sIGSF adhesion complexes CD2-CD58, but their localization around the cSMAC was completely different in a minimal IS in which only the integrin adhesion complex and TCR-pMHC specificity complex were involved (Fig. 2A-first row). While sIGSF localized in the dSMAC, the accumulation of costimulatory complexes around the core of TCR-pMHC complex in the cSMAC was a combined result of SBS, passively following TCR-pMHC, and being actively transported towards the center due to interaction with the F-actin centripetal flow (Fig. 2A-second row).

**Fig. 2.**
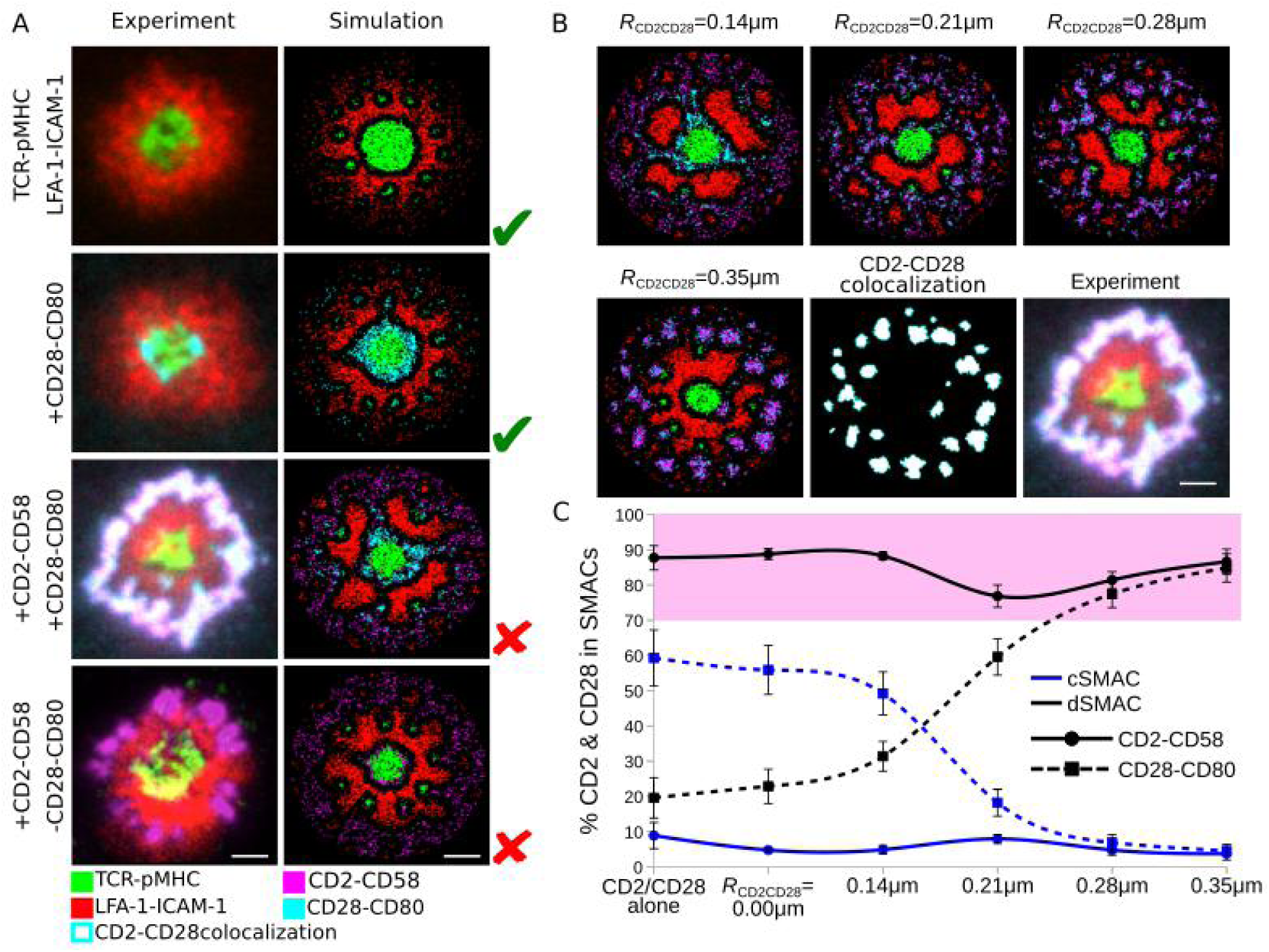
CD28-CD80 complexes relocate to the CD2-CD58 corolla based on an attractive force between both complexes. (**A**) Experimentally observed IS patterns including different receptor-ligand complexes (left column) and comparison with IS patterns obtained by simulations (right column). (**B**) Attraction between CD2-CD58 and CD28-CD80 complexes within radius *R*_CD2CD28_=0.14-0.35μm and weight *W*_CD2CD28_=1.0, colocalization of CD2-CD58 and CD28-CD80 complexes for *R*_CD2CD28_=0.35μm and experimental data. (**C**) Amount of CD2-CD58 (solid) and CD28-CD80 (dashed) complexes in the central and distal SMACs for various CD2-CD28 attractive force radii. Pink shaded area denotes the experimental range of CD2-CD58 and CD28-CD80 in the dSMAC. Parameters from Table 1. Error bars represent SD of N=10 simulations. Scale bar: 2μm. TCR-pMHC: green, LFA-1-ICAM-1: red, CD2-CD58: magenta, CD28-CD80: cyan, CD2-CD28 colocalization: white.

Interestingly, it was recently shown experimentally that the sIGSF adhesion complex can relocate costimulatory/checkpoint complexes to the corolla (Fig. 2A-third row) (*11*). Considering the above-mentioned experimental and theoretical results for CD28 (*16, 19*), we introduced CD2 and CD58 molecules into the model. The association and dissociation rates were taken from the literature (*23-26*). Due to the similar sizes of CD2-CD58, CD28-CD80 and TCR-pMHC complexes (≈12.8-13.1 nm), we did not assume any interaction between them, but only introduced SBS between each of these and LFA-1-ICAM-1 complexes.

**Table 1:** Reference parameter values, used throughout the article.

| Name | Parameter | Value | Reference |
| --- | --- | --- | --- |
| Radius of contact region | $R$ ($\mu\text{m}$) | 4.9 | (17, 54) |
| Lattice constant | $a$ ($\mu\text{m}$) | 0.07 | (17, 54) |
| Free molecule diffusion constant | $D_m$ ($\mu\text{m}^2/\text{s}$) | 0.10 | (17, 54) |
| Complex diffusion constant | $D_c$ ($\mu\text{m}^2/\text{s}$) | 0.06 | (17, 54) |
| Radius of SBS between TCR-pMHC and LFA-1-ICAM-1 | $R_{\text{SBS,TCR-LFA}}$ ($\mu\text{m}$) | 0.42 | (17, 54) |
| Weight of SBS between TCR-pMHC and LFA-1-ICAM-1 | $W_{\text{SBS,TCR-LFA}}$ | -1.0 | (17, 54) |
| Radius of SBS between CD28-CD80 and LFA-1-ICAM-1 | $R_{\text{SBS,CD28-LFA}}$ ($\mu\text{m}$) | 0.42 | (17) |
| Weight of SBS between CD28-CD80 and LFA-1-ICAM-1 | $W_{\text{SBS,CD28-LFA}}$ | -1.0 | (17) |
| Radius of SBS between CD2-CD58 and LFA-1-ICAM-1 | $R_{\text{SBS,CD2-LFA}}$ ($\mu\text{m}$) | 0.42 | fixed |
| Weight of SBS between CD2-CD58 and LFA-1-ICAM-1 | $W_{\text{SBS,CD2-LFA}}$ | -1.0 | fixed |
| Radius of SBS between CD45 and TCR-pMHC | $R_{\text{CD45TCR}}$ ($\mu\text{m}$) | 0.28 | varied:<br>0.00-0.42 |
| Weight of SBS between CD45 and TCR-pMHC | $W_{\text{CD45TCR}}$ | -1.0 | fixed |
| Radius of SBS between CD45 and LFA-1-ICAM-1 | $R_{\text{CD45LFA}}$ ($\mu\text{m}$) | 0.00 | varied:<br>0.00-0.28 |
| Weight of SBS between CD45 and LFA-1-ICAM-1 | $W_{\text{CD45LFA}}$ | 0.0 | varied:<br>-1.00-0.00 |
| Radius of SBS between CD45 and CD2-CD58 | $R_{\text{CD45CD2}}$ ($\mu\text{m}$) | 0.28 | varied:<br>0.21-0.42 |
| Weight of SBS between CD45 and CD2-CD58 | $W_{\text{CD45CD2}}$ | -1.0 | fixed |
| Radius of Attraction between CD2-CD58 and CD28-CD80 | $R_{\text{CD2CD28}}$ ($\mu\text{m}$) | 0.35 | varied:<br>0.00-0.42 |
| Weight of Attraction between CD2-CD58 and CD28-CD80 | $W_{\text{CD2CD28}}$ | 1.0 | fixed |
| Radius of Attraction between CD2-CD58 and CD2-CD58 (Self Attraction) | $R_{\text{SelfAtt}}$ ($\mu\text{m}$) | 0.28 | varied:<br>0.00-0.42 |
| Weight of Attraction between CD2-CD58 and CD2-CD58 (Self Attraction) | $W_{\text{SelfAtt}}$ | 1.0 | varied:<br>0.2-1.0 |
| Dissociation rate TCR-pMHC | $k_{\text{off,TCR}}$ (1/s) | 0.1 | (17, 54) |
| Association rate TCR-pMHC | $k_{\text{on,TCR}}$ (1/Ms) | $2 \cdot 10^4$ | (17, 54) |
| Dissociation rate LFA-1-ICAM-1 | $k_{\text{off,LFA}}$ (1/s) | 0.03 | (17, 54) |
| Association rate LFA-1-ICAM-1 | $k_{\text{on,LFA}}$ (1/Ms) | $3 \cdot 10^5$ | (17, 54) |
| Dissociation rate CD28-CD80 | $k_{\text{off,CD28}}$ (1/s) | 1.6 | (17) |
| Association rate CD28-CD80 | $k_{\text{off,CD28}}$ (1/Ms) | $4 \cdot 10^5$ | (17) |
| Dissociation rate CD2-CD58 | $k_{\text{on,CD2}}$ (1/s) | 4 | (20-23) |
| Association rate CD2-CD58 | $k_{\text{off,CD2}}$ (1/Ms) | $6 \cdot 10^5$ | (20-23) |
| TCR-pMHC centrally directed force | $C_{\text{TCR}}$ | 1.00 | varied:<br>0.00-1.00 |
| LFA-1-ICAM-1 centrally directed force | $C_{\text{LFA}}$ | 0.06 | varied:<br>0.00-0.10 |
| CD28-CD80 centrally directed force | $C_{\text{CD28}}$ | 0.20 | (17) |
| CD2-CD58 centrally directed force | $C_{\text{CD2}}$ | 0.00 | varied:<br>0.00-0.10 |
| Density of TCR, pMHC | (1/$\mu\text{m}^2$) | 9 | varied: 9-14 |
| Density of LFA-1, ICAM-1 | (1/$\mu\text{m}^2$) | 23 | fixed |
| Density of CD28, CD80 | (1/$\mu\text{m}^2$) | 11 | fixed |
| Density of CD2, CD58 | (1/$\mu\text{m}^2$) | 18 | varied:<br>1-100 |
| Density of CD45 | (1/$\mu\text{m}^2$) | 21 | fixed |

Introduction of CD2-CD58 complexes into the synapse did not affect the annular localization of CD28-CD280 costimulatory complexes, in contrast to the experimental observation (Fig. 2A-third row). This was also confirmed by measuring the amount of CD2-CD58 and CD28-CD80 complexes in the cSMAC and dSMAC of the IS (Fig. 2C). The majority of CD28-CD80 (≈60%) still resided in the cSMAC, while almost 90% of CD2-CD58 complexes were excluded towards the dSMAC, where they resided in a ring-like structure.

The exclusion of CD2-CD58 complexes was the combined result of the LFA-1-ICAM-1 gradient and space competition in the cSMAC and pSMAC with complexes that were actively transported towards the center by coupling to the F-actin flow, i.e. TCR-pMHC, LFA-1-ICAM-1 and CD28-CD80 complexes. This set of simulations showed that: (i) in order to relocate costimulatory complexes from the cSMAC to the dSMAC, additional mechanisms are required; and (ii) despite the exclusion of sIGSF adhesion complexes, clustering as found in experiment (see corolla pattern in Fig. 2A-bottom row) did not appear in the dSMAC, also suggesting that additional mechanisms are required. In the following parts of this study we addressed understanding the mechanisms responsible for the relocation of CD28-CD80 to the dSMAC as well as the CD2-CD58 corolla pattern formation.

For the relocation of costimulatory complexes to the dSMAC, the similarities of sIGSF adhesion and costimulatory complexes in protein interaction network and size, could suggest that the two kinds of complexes prefer to stay together during IS formation, as found experimentally (*11*). This was modeled as an attractive interaction defined by interaction radius (*R*_CD2CD28_) and strength (*W*_CD2CD28_). The attractive force was able to overcome the centripetal force of CD28-CD80 and pull them into the dSMAC region (Fig. 2B, C, Movie S1). The greater the attraction radius the more costimulatory complexes relocated in the dSMAC. The amount of CD28-CD80 in the dSMAC increased from ≈20% to ≈80%. The attraction was also sufficient to cluster CD2-CD58 and CD28-CD80 complexes (Fig. 2B), recapitulating the experimentally observed corolla pattern (*11*).

In order to further investigate how the corolla was affected by the LFA-1-ICAM-1 gradient, *in silico* simulations were performed with different strengths of the centripetal force on LFA-1-ICAM-1 complexes (*C*_LFA_=0.00-0.20). The attraction of the CD2-CD58 and CD28-CD80 complexes was kept constant at *R*_CD2CD28_=0.35μm with strength *W*_CD2CD28_=1.0 (Fig. S1). In the absence of centripetal forces (*C*_LFA_=0.00), no LFA-1-ICAM-1 gradient developed and the CD2-CD28 clusters localized to the pSMAC region rather than being excluded to the dSMAC. As the centripetal force increased in strength (*C*_LFA_≥0.03), the CD2-CD28 clusters were excluded to the dSMAC. This result shows the importance of the LFA-1-ICAM-1 gradient for exclusion of sIGSF adhesion and costimulatory complexes to the dSMAC during IS formation.

In contrast to the simulation, CD2-CD58 complexes still formed the corolla pattern on SLB even in the absence of costimulatory complexes (Fig. 2A-bottom row) (*11*). This result suggested that there is one or more missing mechanism in the model responsible for the generation of the corolla pattern. Therefore, we investigated different hypotheses and possible mechanisms.

### sIGSF receptor titration fails to reproduce the corolla pattern

As discussed earlier, the LFA-1-ICAM-1 gradient in the pSMAC together with SBS between LFA-1-ICAM-1 and CD2-CD58 complexes results in the exclusion of CD2-CD58 complexes towards the dSMAC. The *in silico* experiments showed that CD2-CD58 and TCR-pMHC complexes initially colocalize in the microclusters (Fig. 3A). Measuring the amount of CD2-CD58 in each region of the IS (c, p, dSMAC), we observed that during the first two minutes of IS formation the *in silico* and *in vitro* data did not match (Fig. 3B). *In vitro*, this is the spreading phase of the T cell on the SLB, while *in silico* the full IS area is already formed. When this phase is over (≥3 minutes), the *in vitro* and *in silico* data are in good agreement (Fig. 3B), as can also be seen from the radial density profiles at 10 minutes of IS formation (Fig. 3C). Interestingly, a small population of CD2-CD58 complexes passively followed the TCR-pMHC movement toward the center of the IS and localized around the cSMAC (Fig. 3A, B). This population was termed passive followers (*19*). The majority of CD2-CD58 (≈80%), though excluded to the dSMAC, were correctly positioned but failed to form the corolla pattern (Fig. 3A-C).

**Fig. 3.**
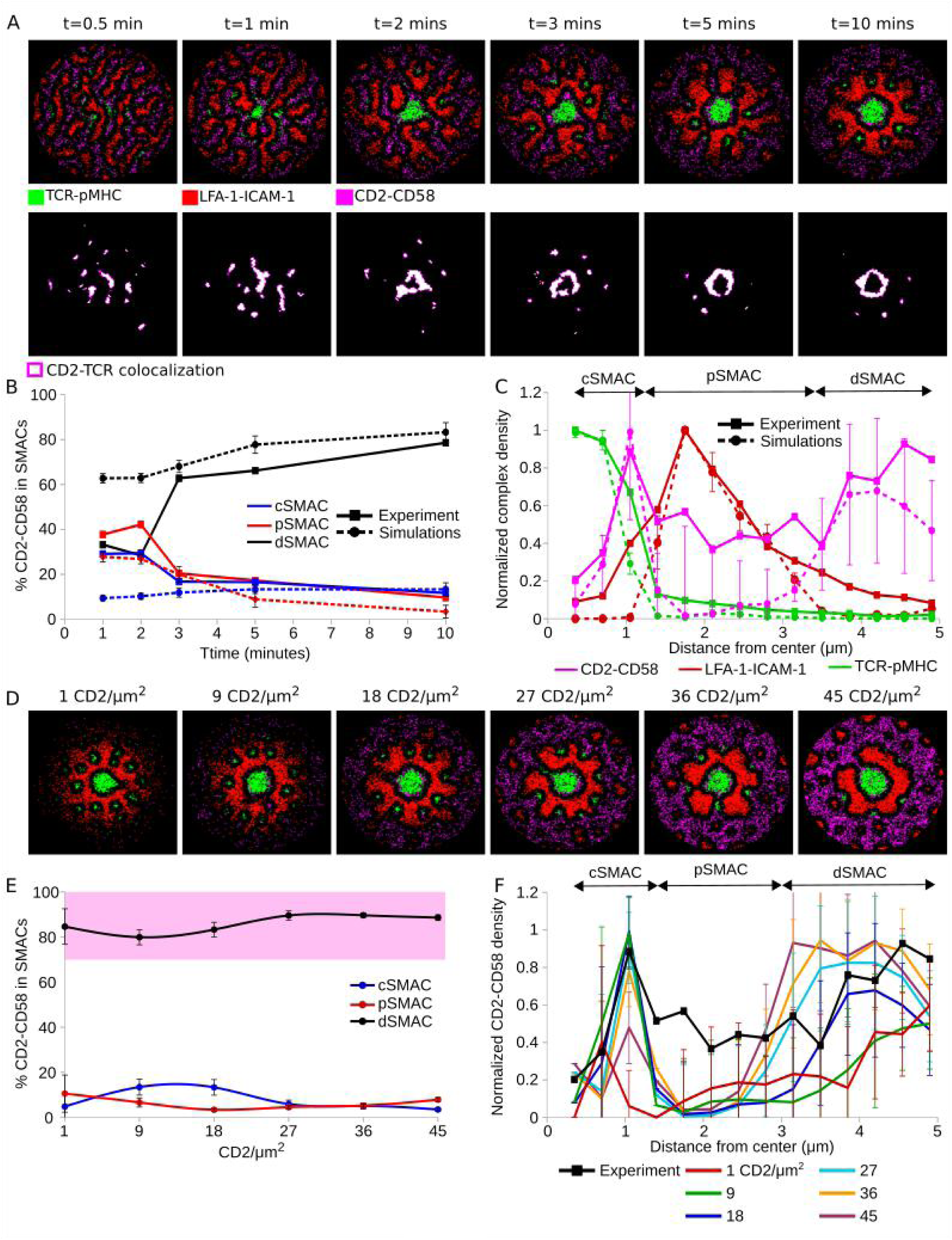
Titration of CD2 in the IS. (**A**) IS formation dynamics between 0.5 and 10 min. (**B**) Amount of CD2-CD58 complexes in central, peripheral and distal SMACs during IS formation in simulation versus experiment (*10*). (**C**) Radial density profile of TCR, LFA-1 and CD2 complexes along the distance from the center at 10 min. (**D**) Titration of the initial CD2 amount from one to 45 molecules/μm^2^. (**E**) Amount of CD2-CD58 complexes in central, peripheral and distal SMACs for different CD2 densities. Pink shaded area denotes the experimental range in the dSMAC. (**F**) Radial density profiles of CD2-CD58 complexes along the distance from the center for different CD2 densities. Parameters from Table 1. Varied parameters: 1-45 CD2/μm^2^; 100 CD58/μm^2^. Error bars represent SD of N=10 simulations. TCR-pMHC: green, LFA-1-ICAM-1: red, CD2-CD58: magenta, CD2-TCR colocalization: white.

Demetriou et al reported that the relative levels of CD2 in different T cell types, including naive and memory CD4+ and CD8+ T cells, and in tumor infiltrating lymphocytes (TILs) of colorectal cancer (CRC) patients versus cells from surrounding healthy tissues might play an important role in the emergence of the corolla pattern (*11*). TILs of CRC patients express smaller numbers of CD2 per T cell, resulting in an impaired or completely absent corolla pattern. Therefore, we sought to investigate whether titration of the CD2 amount would result in the corolla pattern in the dSMAC region.

The amount of CD58 in the SLB lattice was abundant (100 CD58/μm^2^), discounting its role as a limiting factor in CD2-CD58 complex formation. We then studied the localization of CD2-CD58 complexes, starting with population sizes as low as 1 and reaching up to 45 CD2/μm^2^. With increasing CD2 density, the CD2-CD58 ring in the dSMAC became more prominent (Fig. 3D), but the distinct “petals” of the corolla pattern did not emerge. Further, as shown in Fig. 3E increasing CD2 density did not alter the amount of excluded CD2-CD58 (≈90%), reaching the same levels as in Fig. 2C and 3B. Interestingly, as a result of crowding and SBS between CD2-CD58 and LFA-1-ICAM-1, the passive follower population in the outer cSMAC saturated at a maximum capacity already at 18 CD2/μm^2^ (Fig. 3F). The lack of CD2-CD58 complex clustering in the dSMAC suggests that the amount of CD2 or other similar sIGSF family receptor is not a sufficient driver for corolla formation.

### sIGSF adhesion complex self-attraction is sufficient for corolla pattern

We further tested mechanisms able to cluster the excluded CD2-CD58 complexes. Both, centripetal forces weaker or similar to those acting on LFA-1-ICAM-1 (Fig. S2A, B), and adhesion between neighboring CD2-CD58 complexes (Fig. S2C) failed to reproduce clustering in the dSMAC and thus the corolla.

We hypothesized an attractive interaction between neighboring CD2-CD58 complexes. Such an attraction could be the result of CD2 interacting with F-actin asters that are not part of the ramified transport network (*41*). This self-attraction was implemented as an attractive force between CD2-CD58 complexes with varying radii, *R*_SelfAtt_=0.00-0.42μm, and strengths *W*_SelfAtt_ ϵ (0, 1]. For attraction radius *R*_SelfAtt_=0.28 μm and CD2 concentrations ≥18/μm^2^ distinct clustering appeared, emulating the corolla pattern (Fig. 4A). Increased CD2 density on the T cell surface led to a linear increase of the number of cells developing a corolla pattern *in vitro* (*11*). Titration of the CD2 amount with fixed attraction radius *R*_SelfAtt_=0.28 μm divided CD2-CD58 into two populations, one localized around the cSMAC and the other in the corolla (Fig. 4B). With increasing amount of CD2, the sIGSF adhesion complex population in the corolla increased in a linear fashion (see linear regression line for CD2 density ≤4/μm^2^ in Fig. 4B) before saturation at ≈90%, as before (Fig. 2C, 3B, E). Assuming ergodicity, the linear increase of the number of cells with developed corolla *in vitro* (*11*) corroborates the linear increase of the number of CD2-CD58 complexes in the corolla with increasing CD2 density *in silico*.

**Fig. 4.**
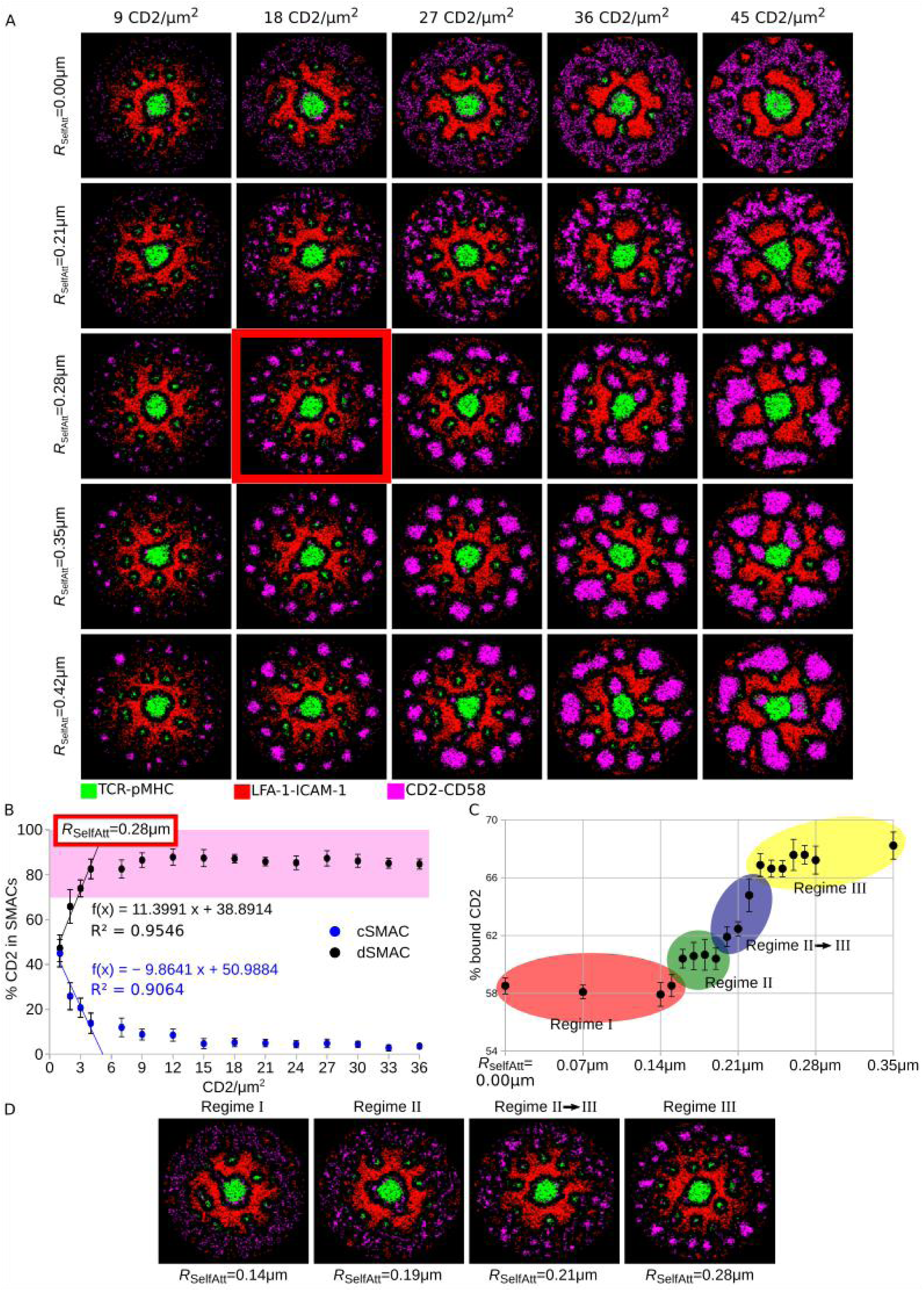
CD2 self-attraction results in the corolla pattern. (**A**) Rows: Different radii of self-attraction, *R*_SelfAtt_, between CD2-CD58 complexes. Columns: Different densities of CD2 molecules. (**B**) Amount of CD2-CD58 complexes in the central and distal SMAC. Pink shaded area denotes the experimental range in the dSMAC. The function *f*(x) is a linear regression based on the four lowest CD2 densities, i.e. ≤ 4/μm^2^. (**C, D**) Amount of CD2 that formed complexes with CD58 and the emergence of four regimes based on the amount of CD2-CD58 complex formation and the clustering in the IS. Parameters from Table 1. Varied parameters: 1-45 CD2/μm^2^; 100 CD58/μm^2^. Error bars represent SD of N=10 simulations. TCR-pMHC: green, LFA-1-ICAM-1: red, CD2-CD58: magenta.

Interestingly, stronger self-attraction increased the amount of CD2 molecules that formed complexes with CD58 (Fig. 4C). Comparing the amount of bound CD2 with the state of clustering in the dSMAC, we identified four regimes (Fig. 4C, D). In Regime I, *R*_SelfAtt_≤0.15μm, CD2-CD58 were located in the dSMAC but not clustered. In Regime II, 0.16≤*R*_SelfAtt_≤0.19μm, some individual CD2-CD58 stuck together while in Regime II→III, a transition regime, 0.20≤*R*_SelfAtt_≤0.22μm, denser CD2-CD58 areas appeared. Finally, in Regime III, *R*_SelfAt_≥0.23μm the CD2-CD58 clustering was more pronounced and the corolla pattern became apparent. It is remarkable that the simulations suggest a sharp transition from regime II to III.

For the case of self-attraction with radius *R*_SelfAtt_=0.28μm, we observed that CD2-CD58 complexes initially (t<3 minutes) colocalized with TCR-pMHC in the microclusters (Fig. S3 (top and middle row) and Movie S2). As TCR-pMHC accumulated in the cSMAC (t>3 minutes), some of the CD2-CD58 complexes passively followed while the majority were relocated in the dSMAC, where the corolla pattern eventually formed (t=10 minutes). This recapitulates the path described in experiment (*11*). Plotting the tracks of twenty random CD2-CD58 complexes (Fig. S3 (bottom row) and Movie S3) we saw that they initially moved inwards and when the LFA-1-ICAM-1 complexes formed in the pSMAC they were excluded towards the dSMAC. The strength of the self-attraction, *W*_SelfAtt_ ϵ (0, 1], did not alter these results (Fig. S4): The transition from the CD2-CD58 ring in the dSMAC to the corolla occurred for *R*_SelfAtt_>0.21μm (Fig. S4A), while their localization in the IS was not substantially affected (Fig. S4B-D).

These results together showed that a phenomenological attractive force between sIGSF adhesion complexes, possibly originating from their interaction with local F-actin asters or networks of signaling proteins like Lck, can reproduce the experimentally observed corolla pattern.

### Glycocalyx proteins residing in the dSMAC promote corolla pattern

In reality many more molecules are present in the IS region contributing to a dense glycocalyx. Next, we asked whether such molecules affect the sIGSF adhesion complex localization and the transition from the ring in the dSMAC to the corolla pattern. An example is the transmembrane protein tyrosine phosphatase CD45. CD45 molecules are important for T cell activation (*34-38*) and their concentration in the dSMAC (*35, 39, 40*) positions them to regulate sIGSF adhesion complexes in this compartment.

At first, we asked how CD45 molecules are excluded from the IS. Their size, ≈20nm, is smaller than the maximal reach of LFA-1-ICAM-1 complexes and the exclusion from the TCR-pMHC microclusters (*35, 40*) suggested SBS between CD45 and TCR-pMHC. For simplicity, we first investigated the IS dynamics only with TCR, LFA-1 and CD45 on the T cell and pMHC and ICAM-1 on the SLB. The repulsive force between TCR-pMHC and CD45 with radius *R*_CD45TCR_ led to initial exclusion of CD45 molecules from TCR microclusters. Together with the LFA-1-ICAM-1 gradient, CD45 was eventually excluded from the cSMAC and inner pSMAC, restricting CD45 to the outer pSMAC and dSMAC (Fig. S5A, C). Accordingly, we investigated how additional SBS between LFA-1-ICAM-1 complexes and CD45 affects the IS dynamics and pattern. A very weak repulsion, i.e. small radius of repulsion (*R*_CD45LFA_=0.14μm), between the LFA-1-ICAM-1 complexes and CD45 resulted in complete exclusion of CD45 from the pSMAC. Interestingly, for increasing radii, *R*_CD45LFA_>0.21μm, clusters of CD45 appeared in the pSMAC and around the cSMAC (Fig. S5B). This resulted in a slightly decreasing amount of CD45 in the dSMAC, but still more than 80% remained there (Fig. S5D). Since CD45 is experimentally shown to form a ring in the dSMAC (*35, 40*), we decided to include SBS only between TCR-pMHC and CD45.

TCR-pMHC and CD2-CD58 complexes have similar sizes and therefore we included the same SBS between both of these and CD45. No self-attraction between CD2-CD58 was included in these *in silico* experiments. For a repulsion radius of *R*_CD45TCR_=*R*_CD45CD2_=0.21μm, CD2-CD58 complexes colocalized with CD45 in the dSMAC but did not form a distinct corolla (Fig. 5A). The corolla emerged for *R*_CD45TCR_=*R*_CD45CD2_≥0.28μm (Movie S4). Around 80% of the total CD2-CD58 participated in the formation of the corolla in the dSMAC (Fig. 5B), a fraction in accordance with the experimental data (*11*). Interestingly, with increasing repulsion strength, the amount of CD2 molecules that formed complexes with CD58 increased in a linear fashion (Fig. 5C). These results suggested that the corolla pattern formation can also be driven by a phase separation process with competition for space between a percolating phase enriched in CD45 and microdomains enriched in CD2-CD58. Other glycocalyx components could contribute to corolla formation (*42*).

**Fig. 5.**
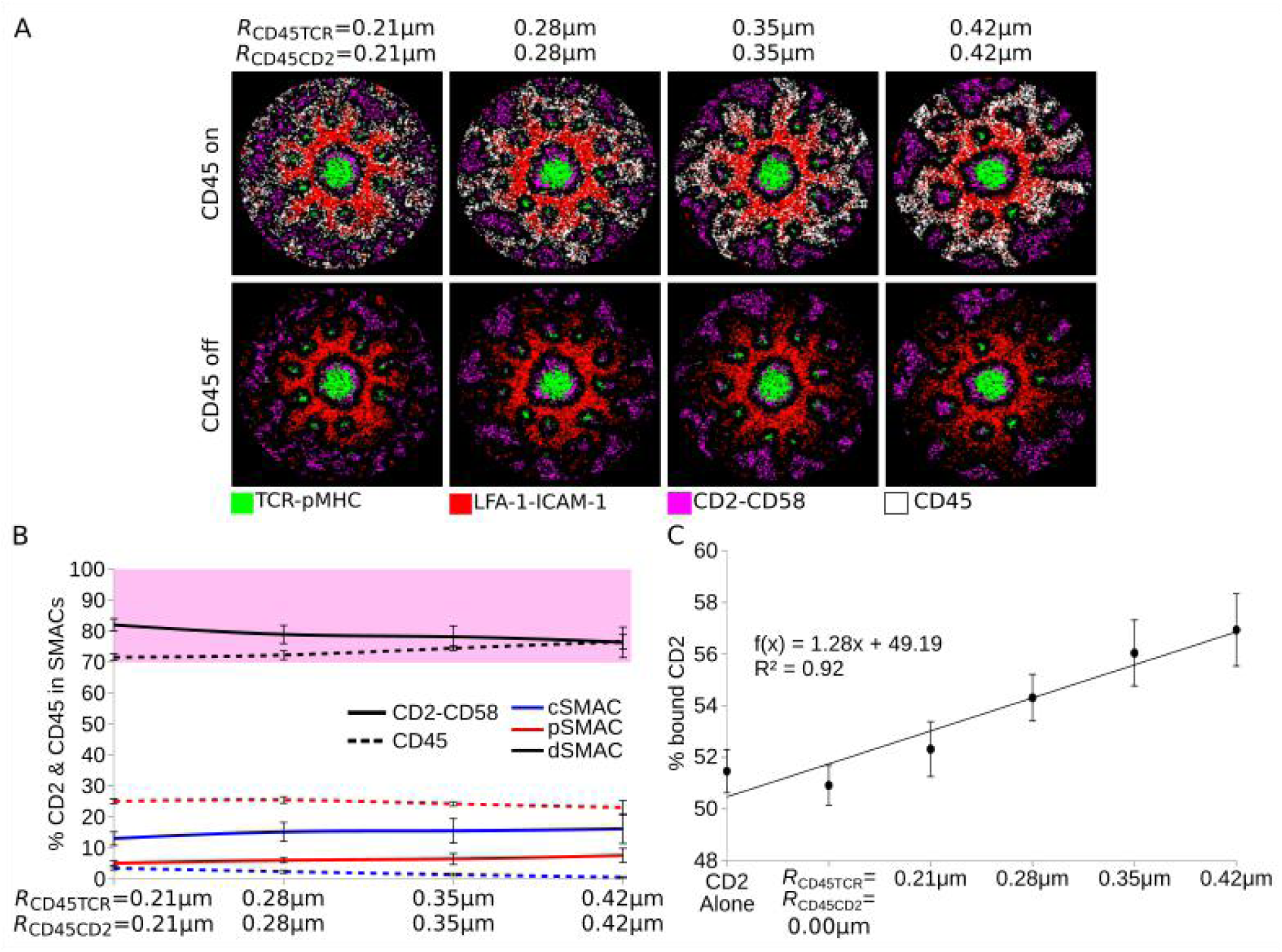
Interaction of CD45 with CD2-CD58 and TCR-pMHC complexes. (**A**) IS pattern at 10 minutes for different radii of repulsion between CD45 and CD2-CD58 as well as CD45 and TCR-pMHC. The same plots are shown with and without resolving CD45. (**B**) Amount of CD45 and CD2-CD58 in the central, peripheral and distal SMACs for different radii of repulsion. Pink shaded area denotes the experimental range of CD2-CD58 in the dSMAC. (**C**) Percent of CD2 molecules that formed complexes with CD58. The function *f*(x) is a linear regression. Parameters from Table 1. Error bars represent SD of N=10 simulations. TCR-pMHC: green, LFA-1-ICAM-1: red, CD2-CD58: magenta, CD45: white.

Additionally, we performed CD2 titration while keeping the repulsion at *R*_CD45TCR_=*R*_CD45CD2_≥0.28μm. For CD2 densities between 1-4/μm^2^, no clustering appeared in the corolla and a large fraction of CD2-CD58 complexes resided in the central IS region (Fig 6A, B). As the CD2 amount increased, not only the majority of CD2 was excluded, reaching more than 70% in the dSMAC, in accordance with the experiment (*11*), but also clustering in the corolla was induced (Fig. 6A, B). These results together showed that the amount of CD2 can affect corolla formation (*11*), provided the presence of large, SBS inducing glycocalyx molecules in the dSMAC. Therefore, reduced CD2 numbers on T cells of CRC TILs, indeed, can result in absence of CD2-CD58 corolla formation, with implications for the balance of costimulatory and checkpoint function, as suggested before (11).

**Fig. 6.**
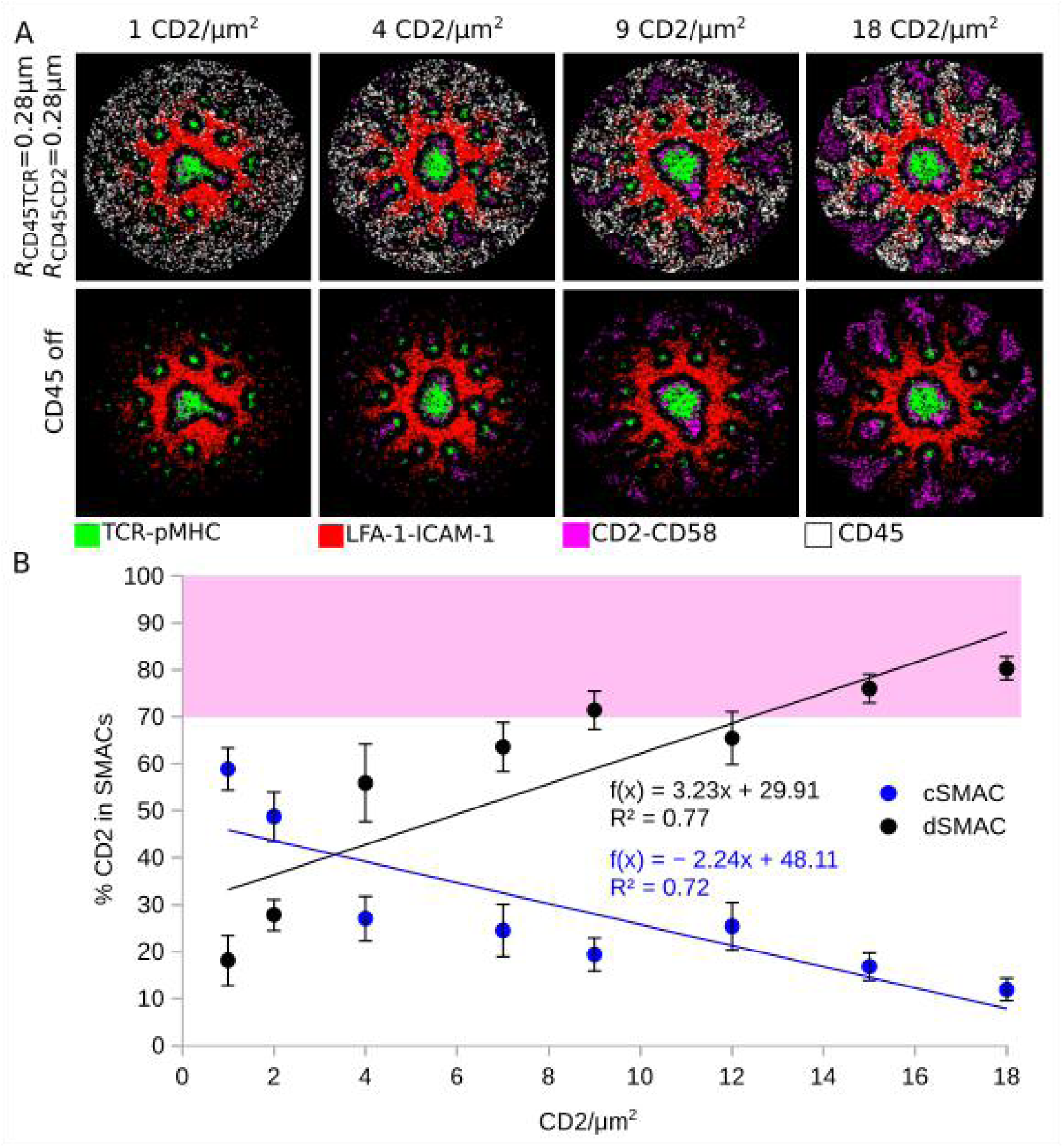
CD2 titration in the presence of CD45. (**A**) Titration of CD2 amount, with constant repulsion between CD45 and CD2-CD58 and TCR-pMHC (*R*_CD45TCR_=*R*_CD45CD2_=0.28μm) with CD45 shown (top row) or not (bottom row). (**B**) Amount of CD2-CD58 in cSMAC and dSMAC, during CD2 amount titration. Pink shaded area denotes the experimental range of CD2-CD58 in the dSMAC. The function *f*(x) is a linear regression. Parameters from Table 1. Varied parameters: 1-18 CD2/μm^2^. Error bars represent SD of N=10 simulations. TCR-pMHC: green, LFA-1-ICAM-1: red, CD2-CD58: magenta, CD45: white.

Finally, in order to confirm the presence and also to show the importance of the LFA-1 gradient, we performed *in vitro* experiments where after allowing the synapse to form for 20 minutes, the cells were treated with Latrunculin A (LatA). LatA is known to disrupt the ramified F-actin transport network, and therefore the centripetal flow of LFA-1 complexes. In accordance to the experiments, we performed simulations where the centripetal flow of LFA-1 (*C*_LFA_) was altered after 20 minutes of synapse formation. Interestingly, we observed both *in vitro* and *in silico*, that depending on the LatA dose, and accordingly the *C*_LFA_ strength, the corolla pattern is disrupted and eventually ceases to exist. CD2-CD58 complexes then move towards the center of the IS (Fig. 7). For LatA=330nM the corolla completely disappeared, as was also observed for *C*_LFA_=0.00 (Fig. 7A). The radial density profiles of CD2 and LFA-1 complexes were similar between experiments and simulations (Fig. 7B). These data provide an additional qualitative validation of the IS simulation model.

**Fig. 7.**
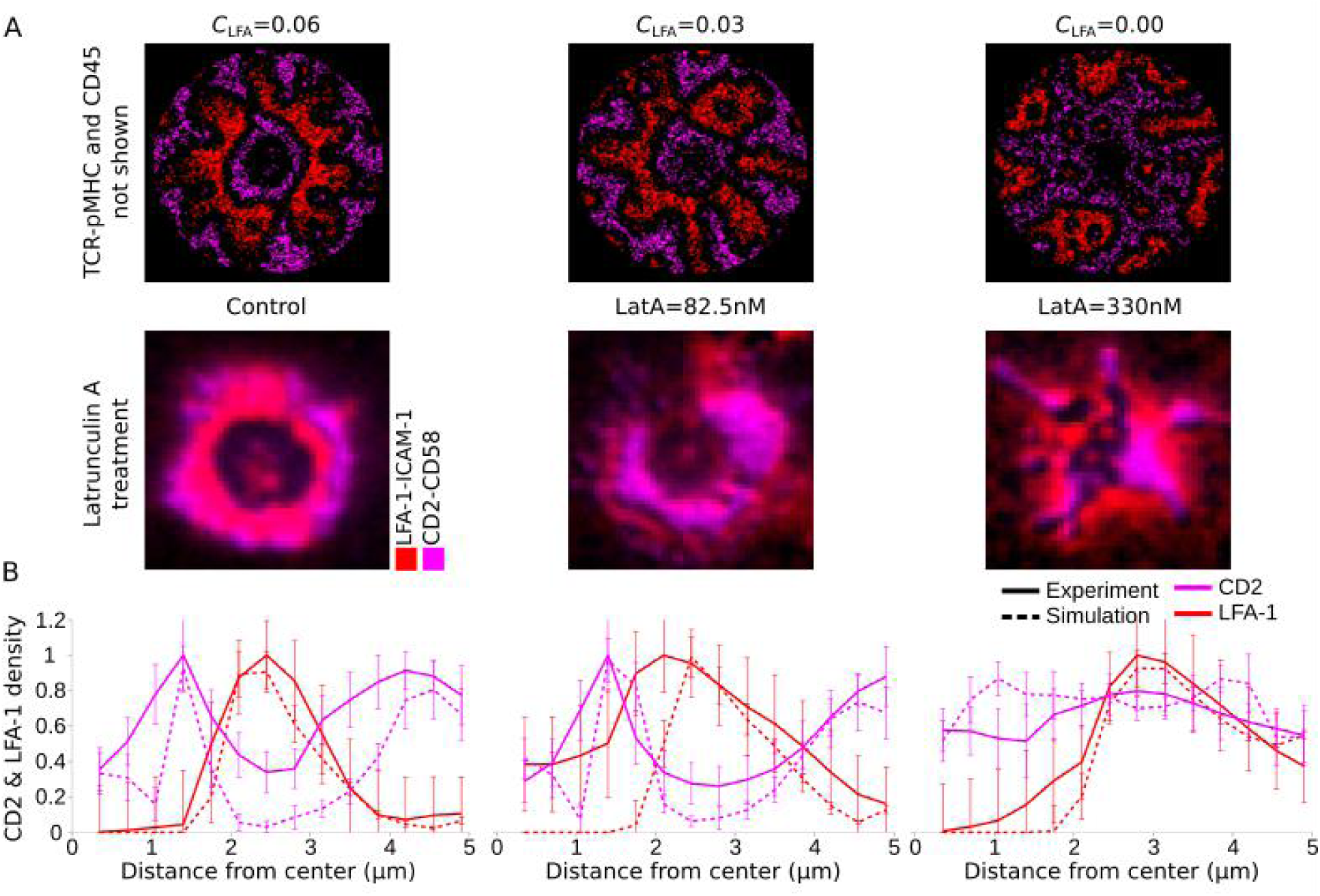
LatA treatment and alteration of the LFA-1 centripetal transport strength (*C*_LFA_), 20 minutes after synapse formation. (**A**) IS simulations of altered *C*_LFA_ strength (top) and cells treated with different doses of LatA. (**B**) Radial density profiles of CD2-CD58 and LFA-1-ICAM-1 complexes along the distance from the center for different *C*_LFA_ strengths (dashed lines) and LatA doses (solid lines). Parameters from Table 1. Varied parameters: 14 TCR, pMHC/μm^2^; 23 CD2, CD58/μm^2^; *C*_LFA_=0.00-0.06. Error bars represent SD of N=10 simulations and at least N=15 treated cells. LFA-1-ICAM-1: red, CD2-CD58: magenta.

## Discussion

A limitation of previous immune synapse models was a failure to incorporate two major classes of molecules. Our studies and others have simulated basic features of IS formation related to the cSMAC and pSMAC (*43-46*). Recently, we refined the model to incorporate costimulation complexes and a more accurate simulation of the pSMAC (*19*). We also now recognize that rules developed for costimulatory interactions likely also apply to checkpoint molecules that counteract costimulatory signals (*3*). In this study we have generalized this agent-based model to incorporate sIGSF adhesion complexes and large glycocalyx molecules like CD45 in order to simulate a newly described feature of human T cell IS (*11*). The *in silico* experiments pointed to two possible mechanisms contributing to corolla formation. One active, energy consuming mechanism, where self-attraction between sIGSF adhesion complexes led to their clustering and together with exclusion by the integrin adhesion complex gradient, to the formation of the corolla pattern (Fig. 4, S3). The CD2 corolla can also be formed by an extracellular size dependent phase separation mechanism in combination with competition for space. In this case, the presence of other, SBS inducing molecules in the dSMAC, such as glycocalyx component CD45, led to sIGSF adhesion complex clustering and consequently the corolla pattern (Fig. 5, 6). These molecular and mechanistic features establish a general model and simulation platform that can recapitulate complex pattern formation processes observed in cell-bilayer and cell-cell interfaces.

We have categorized many potential interactions into larger groups based on specificity (TCR-pMHC), integrin adhesion complexes (LFA-1-ICAM-1), sIGSF adhesion complexes (CD2-CD58) and costimulatory/checkpoint complexes (CD28-CD80 and PD-1-PDL1). We expected that methods we have used to simulate sIGSF adhesion, also apply to hemophilic SLAM family members and heterophilic, but promiscuous adhesion molecules like CR-TAM-CADM1 (*47*). Costimulatory interactions may also include ICOS-ICOSL interactions and tumour necrosis factor (TNF) receptor superfamily members that can interact with sIGSF or TNF superfamily ligands (*11, 48*). The similarities in size and interaction networks of CD2-CD58 sIGSF adhesion complex and CD28-CD80 costimulatory complex together with the indirect links with F-actin (*49*), supported the notion that sIGSF adhesion complexes bind to the centripetal flow of F-actin, just like CD28-CD80 complexes (*19*). Interactions of individual engaged CD2 with F-actin would be transient due to the brief lifetime of CD2-CD58 interactions. These interactions may not result in domain transport as high lateral mobility within the microdomains might instead generate convective mixing within the microdomains and deformation away from perfect circles dictated by self-attraction or size dependent phase separation mechanisms. Although in some cases, a weak centripetal force seemed compatible with corolla formation, the amount of CD2-CD58 complexes residing around the cSMAC contradicted the experimental data (Fig. S2A, B) (*11*). The localization of sIGSF adhesion complexes then more resembled that of costimulatory complexes (*16, 19*), rather than the corolla (*11*). In general, adhesion molecules are expressed at 10-fold greater abundance compared to costimulatory complexes, but our results suggest there are additional differences.

The attractive force between sIGSF adhesion complexes and costimulatory/checkpoint complexes or among sIGSF adhesion complexes that are sufficient for corolla formation in simulation may arise from different types of lateral interactions. There are two classes of interactions to consider: active solids like F-actin asters (*49*) and active liquids such as cytoplasmic condensates implication in TCR signaling (*30*). Linking to an active solid like F-actin cytoskeleton requires adapter proteins. For example, the cytoplasmic domain of CD2 interacts with CapZ through CD2AP (CD2-associated protein) (*49*). This interaction is believed to orchestrate protein patterning in the IS (*8*). The multiple poly-proline motifs in the long, unstructured cytoplasmic domain of CD2 is a signature of proteins that can participate in condensate formation (*50*). There are several possibilities for the implementation of such an interaction including hybrids in which liquid condensates interact with F-actin filaments. The necessary range of self-attraction suggests that the organizing structure would be on the order of >200-350nm. Alternately, short range interactions of neighboring complexes, similar to TCR cross-linking (*51, 52*), was unable to reproduce the CD2-CD58 corolla pattern (Fig. S2C).

The second mechanism able to recapitulate the corolla was a passive mechanism based on SBS and local crowding. CD45 is abundantly expressed on T cells and more generally on all leukocytes, and is important for signaling (*34, 53*). CD45 localization in the dSMAC together with the strong SBS (*35, 40*), suggested a repulsive interaction between the two. The confining effects of this force on sIGSF adhesion complexes may account for corolla formation. In general, the model suggests that crowding in the IS and specifically in the dSMAC can induce pattern formation without self-attraction between the sIGSF adhesion complexes. Additional experiments are required for an improved understanding of whether CD45 or any other kind of molecule or complex present in the dSMAC is responsible for the CD2 corolla formation. A combination of knockouts targeting particular core proteins and glycosylation pathways might be able to sufficiently modulate the glycocalyx of T cells to test this mechanism.

The incorporation of the sIGSF adhesion complex corolla and alternative models for how the structure can form greatly increases the fundamental and practical utility of IS simulations. Genetic variants of CD2 and its ligand CD58 are important in Rheumatoid Arthritis (5*4*). Individuals with variants of CD58 that decreased expression are at greater risk of developing multiple sclerosis (*55*). In cancer, loss of CD58 is associated with relapse in Hodgkins disease (*56*) and low CD2 expression is associated with poor outcome on melanoma (*57*). Moreover, exhausted T cells with low CD2 are observed in CRC patients and has a similar impact on TCR signaling to checkpoint PD-1 engagement (*11*). This general model is now also likely to be predictive for patterns of checkpoint complex forming including interactions of PD-1 and its ligands. Additional features such as cis interaction of CD80 and PD-L1, and PD-1 and PD-L1 between adjacent molecules in the same membrane (*58, 59*) could be rapidly implemented.

## Materials and Methods

### Study Design

This study was designed to examine possible mechanisms leading to characteristic pattern formation in immunological synapses. It confirms the utility of computer simulation in rapidly providing new insights into recently described features of synapses and opens avenues for further investigation of their function in health and disease, aiming to aid the search for novel therapeutic approaches.

### Agent-based model

The agent-based model used in this article was formulated in order to replicate experiments performed on supported lipid bilayers (SLBs) instead of APCs (*19, 60*). The model consists of two square lattices, with each node occupied by only one freely diffusing agent, namely the *in silico* molecules. Each agent can diffuse to one of its eight neighboring nodes (Moore neighborhood). Diffusion is implemented as a random walk with a probability defined by the speed of diffusion and the simulation timestep. Binding and unbinding of molecules can happen only with ligands on the exact same position on the opposite lattices, with probabilities defined by (*61*):

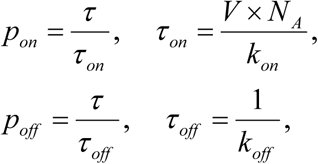

where τ is the simulation timestep, *k*_on_ and *k*_off_ are the on- and off-rates, respectively, V the volume of the complex about to form and *N*_A_ the Avogadro’s number. The *k*_on_ and *k*_off_ for each specific complex forming in the IS are shown in Table 1.

The forces between interacting complexes are modeled as weighted vectors from one complex towards all its interacting neighbors. The forces are exerted within a radius *R*_force_ and all the vectors are summed. The strength of the force is represented as a weight, *W*_force_, where negative weight represents repulsion (SBS) and positive attraction (*19*). In a similar manner, interaction of the IS molecules with the centripetal flow of F-actin is modeled as an empirical centrally directed force added to the vector of the interaction forces discussed above. The strength of this force, *C*_X_ > 0, where X represents each specific complex, shows the coupling strength to F-actin (*19*).

Adhesion between neighboring CD2 pairs was implemented as a phenomenological factor, *g*(*N*), reducing the movement probability, *p*_move_ → *p*_move_*g*(*N*), with *g*(*N*) given by (*54*):

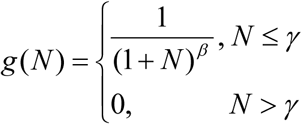

where *β* ϵ [1, 2] is the strength and *γ* ϵ [0, 8] is the number of neighbors considered.

The *in silico* radial density plots are calculated inside equally spaced concentric rings as the fraction of occupied grid points by each type of complex and is normalized by the maximum value. The *in vitro* radial density plots are obtained by measuring the relative fluorescence intensity in 360 line plots across the diameter of the IS and is normalized by the maximum value. The central, peripheral and distal SMACs are defined at the end of the *in silico* experiment, which also determines the amount of each complex in each SMAC in the simulations.

### Statistics

The presented curves and error bars represent the mean values and standard deviation (SD), respectively, of at least ten computer simulations.

## Supplementary Materials

Fig. S1. Effect of the LFA-1 centripetal force, *C*_LFA_, and the emerging pSMAC gradient on the localization of CD2-CD28 clusters.

Fig. S2. Centripetal flow and adhesion between neighboring CD2-CD58 complexes.

Fig. S3. Dynamics of the CD2 corolla pattern formation.

Fig. S4. Alterations in strength, *W*_SelfAtt_, and radii, *R*_SelfAtt_, of the attractive force, with initial CD2 concentration 36 CD2/μm^2^.

Fig. S5. CD45 localization during interaction with TCR-pMHC and LFA-1-ICAM-1 complexes at 10 minutes of IS formation.

Movie S1. CD2-CD28 corolla in the dSMAC for *R*_CD2CD28_=0.35μm, showing 30 minutes of IS formation.

Movie S2. Corolla pattern formation dynamics for *R*_SelfAtt_=0.28μm, showing 30 minutes of IS formation.

Movie S3. Tracks of twenty randomly selected CD2-CD58 complexes during corolla pattern formation *R*_SelfAtt_=0.28μm, showing 30 minutes of IS formation.

Movie S4. Corolla pattern formation due to repulsive interaction between CD45 and CD2-CD58 complexes, for *R*_CD45CD2_=0.28μm, showing 30 minutes of IS formation.

## Author contributions

A.S. conceived and designed the study. A.S. designed the code, performed and analysed the simulations. P.A.R. and M.M.-H. supervised the work. A.S., M.L.D and M.M.-H. wrote and edited the manuscript. P.D., A.K., S.V., V.M. and M.L.D. provided the experimental figures and made intellectual contributions.

## Data and materials availability

The code is written in *C*++ according to all the algorithmic details explained in (*19, 61*), and is available upon request.

