## Supplementary figures for "*In silico* characterization of mechanisms positioning costimulatory and checkpoint complexes in immune synapses"

**Affiliations:**

[#] Current address: Department of Immunology, University of Oslo, Oslo, 0372 Norway.

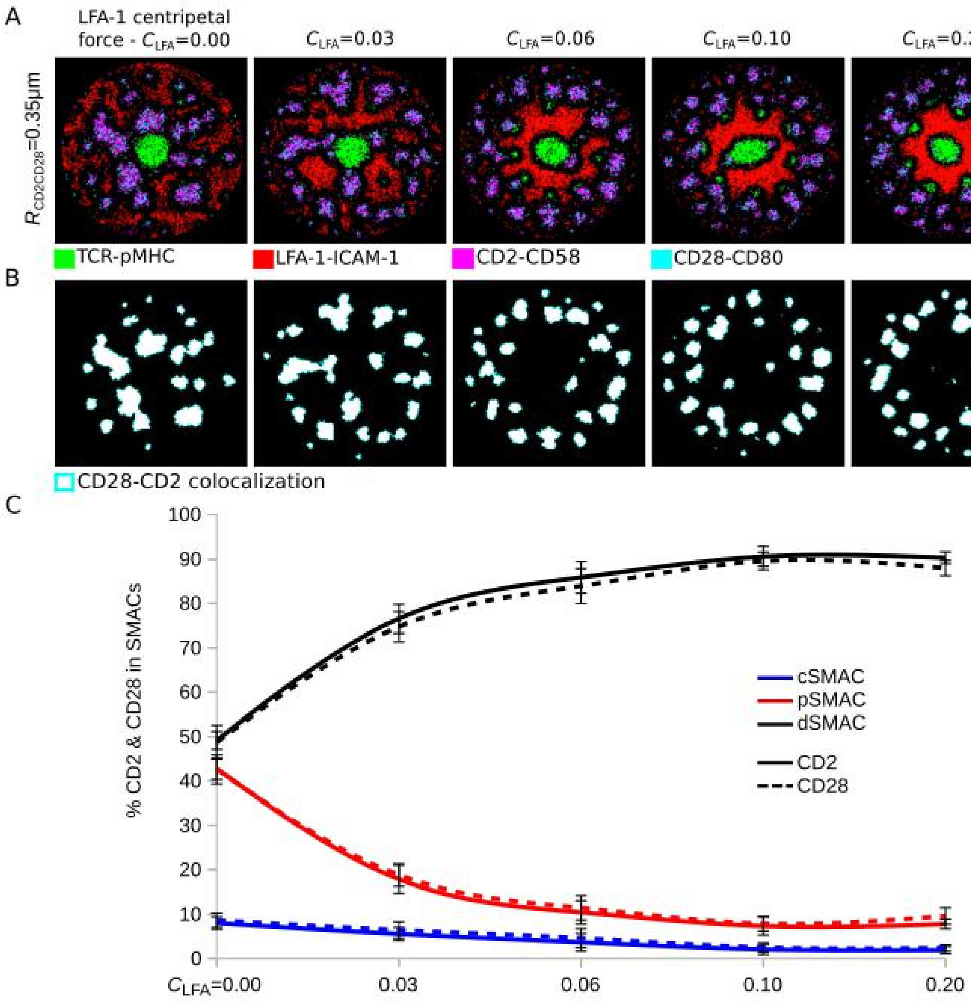


**Fig. S1.** Effect of the LFA-1 centripetal force, $C_{LFA}$, and the emerging pSMAC gradient on the localization of CD2-CD28 clusters. Attraction between CD2-CD58 and CD28-CD80 complexes was $R_{CD2CD28}$=0.35μm and $W_{CD2CD28}$=1.0. (**A**) IS pattern at 10 min. (**B**) Colocalization of CD2-CD58 and CD28-CD80 complexes in the IS at 10 min. (**C**) Amount of CD2-CD58 and CD28-CD80 complexes in different synapse locations in dependence on LFA-1 centripetal force. Parameters from Table 1. Varied parameters: $C_{LFA}$=0.00-0.10. Error bars represent SD of N=10 simulations. TCR-pMHC: green, LFA-1-ICAM-1: red, CD2-CD58: magenta, CD28-CD80: cyan, CD28-CD2 colocalization: white.

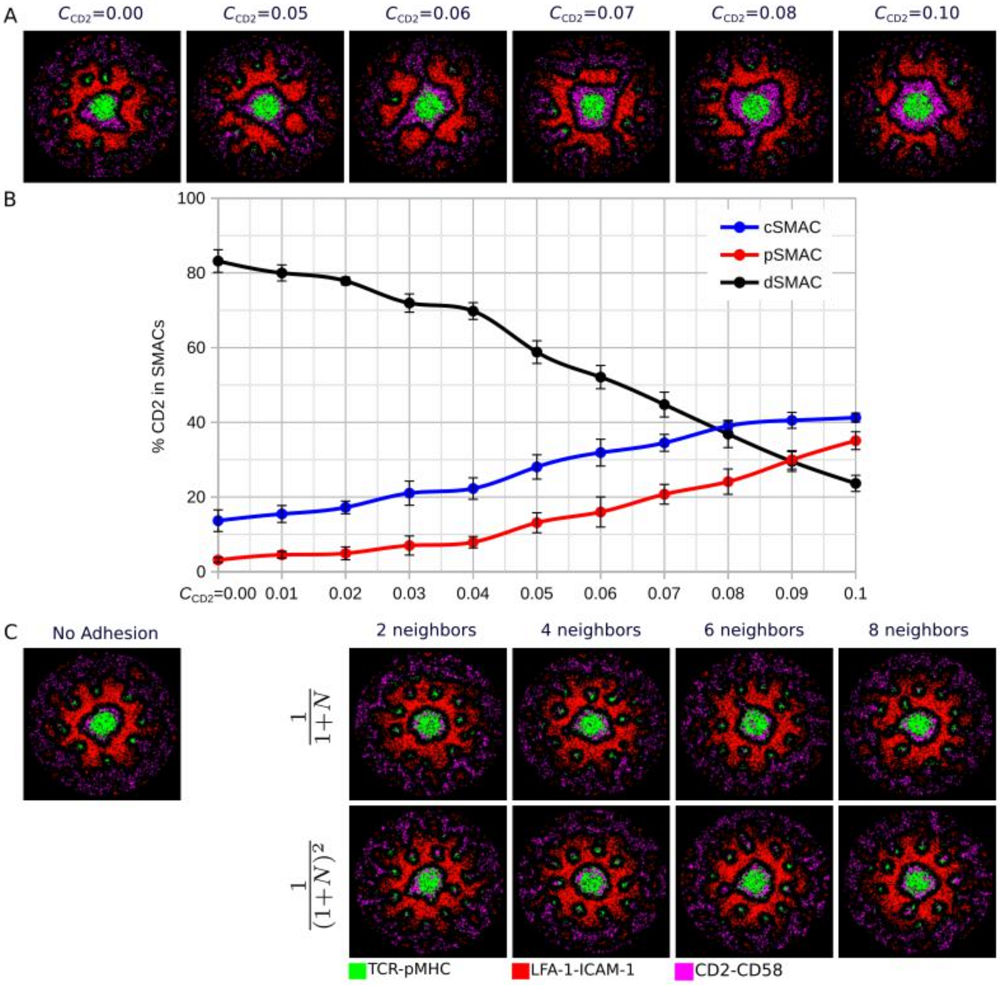


**Fig. S2.** Centripetal flow and adhesion between neighboring CD2-CD58 complexes. (**A**) Centripetal force, $C_{CD2}$, acting on CD2-CD58 complexes. (**B**) Amount of CD2-CD58 in central, peripheral and distal SMACs in dependence on the strength of the centripetal force $C_{CD2}$. (**C**) Adhesion between neighboring CD2-CD58 complexes, given by $g(N)$ (see Methods). Parameters from Table 1. Error bars represent SD of N=10 simulations. TCR-pMHC: green, LFA-1-ICAM-1: red, CD2-CD58: magenta.

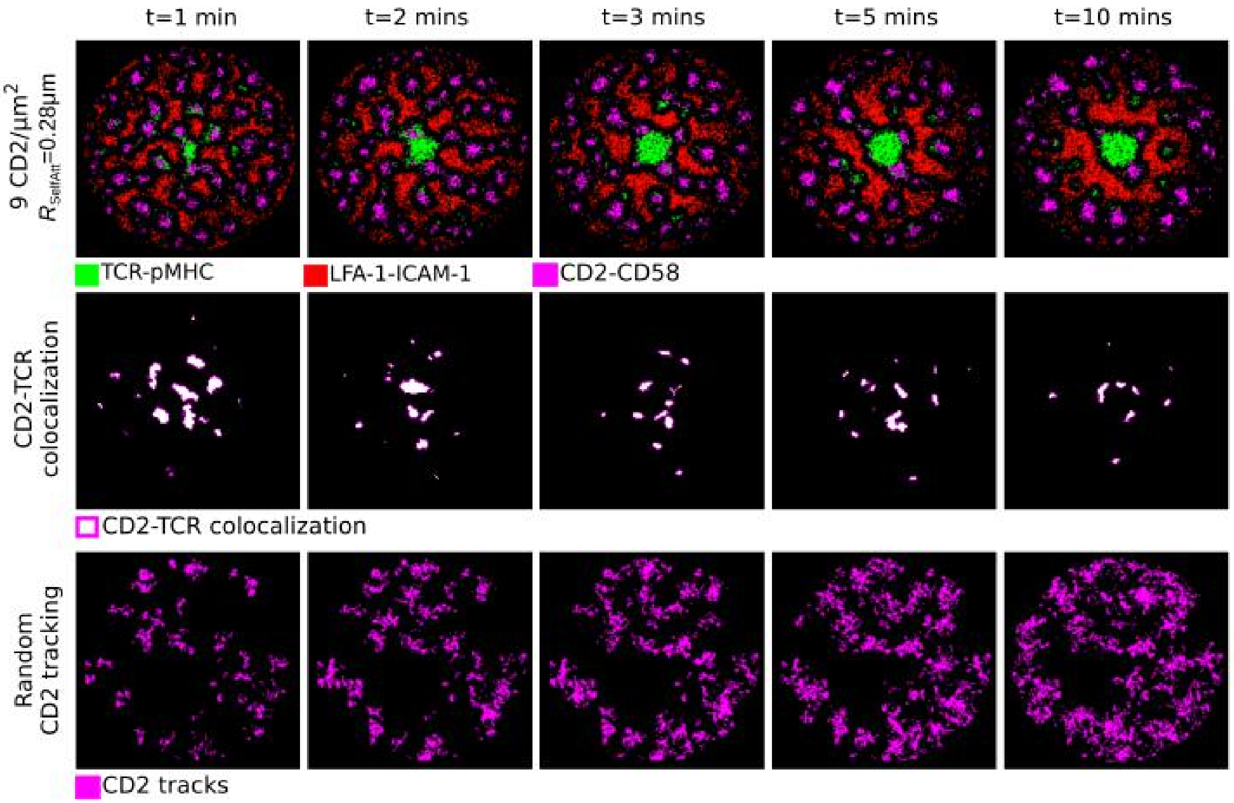


**Fig. S3.** Dynamics of the CD2-CD58 corolla pattern formation together with the colocalization of TCR-pMHC and CD2-CD58 complexes as well as the tracks for twenty randomly selected CD2-CD58 complexes. Parameters from Table 1. $R_{SelfAtt}$=0.28μm, TCR-pMHC: green, LFA-1-ICAM-1: red, CD2-CD58: magenta.

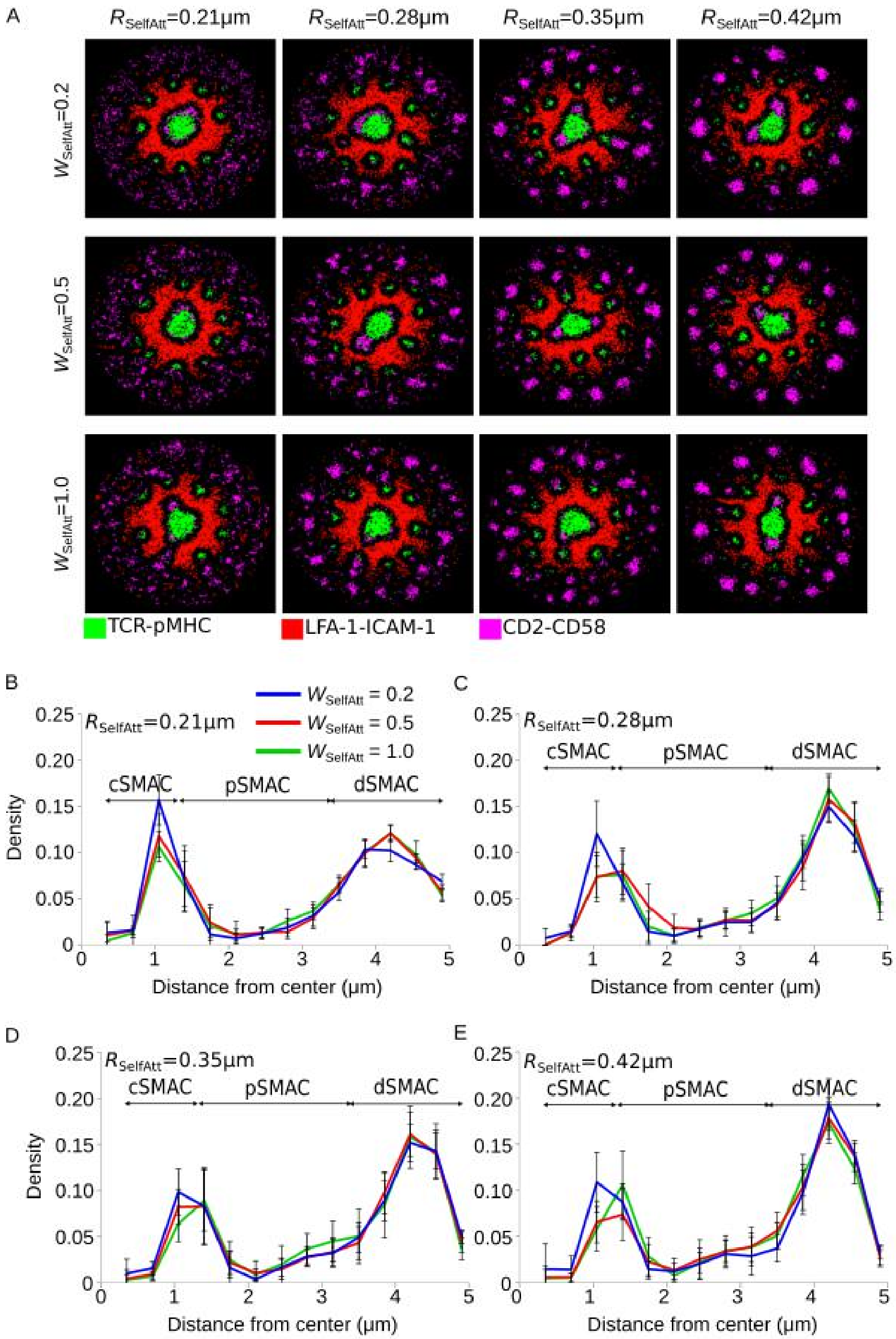


**Fig. S4.** Alterations in strength, $W_{SelfAtt}$, and radii, $R_{SelfAtt}$, of the attractive force between CD2-CD58 complexes, with CD2 density 36 CD2/μm$^2$. (**A** Rows) Different radii of attractive interaction, $R_{SelfAtt}$. (**A** Columns) Different strengths of attractive interaction, $W_{SelfAtt}$. (**B-E**) Radial density profiles of CD2-CD58 complexes along the distance from the center of each column of (**A**). Parameters from Table 1. Varied parameters: $R_{SelfAtt}$=0.21-0.42μm; $W_{SelfAtt}$=0.2-1.0. Error bars represent SD of N=10 simulations. TCR-pMHC: green, LFA-1-ICAM-1: red, CD2-CD58: magenta.

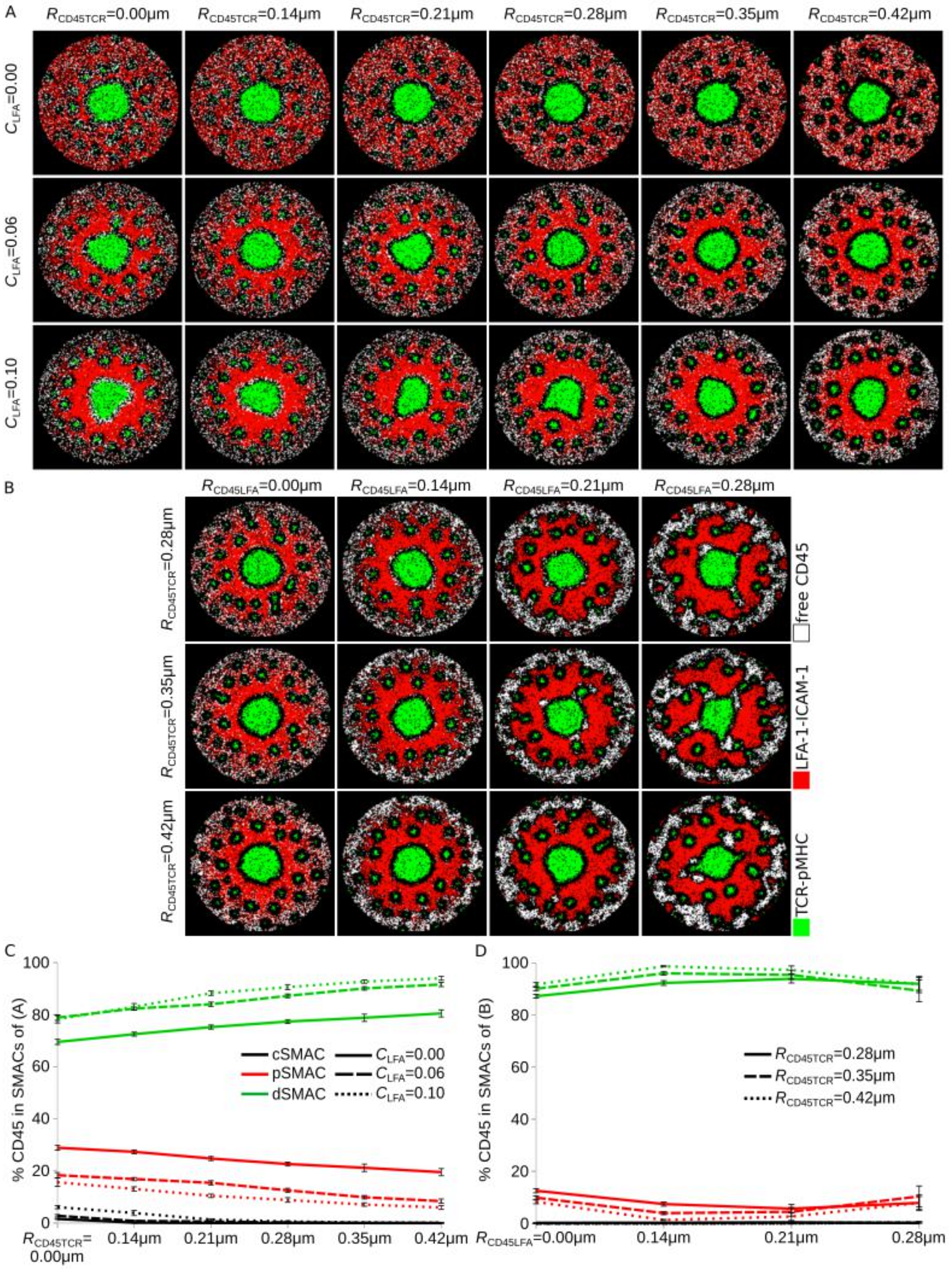


**Fig. S5.** CD45 localization during interaction with TCR-pMHC and LFA-1-ICAM-1 complexes at 10 minutes of IS formation. (**A**) Different centripetal forces of LFA-1-ICAM-1 complexes, $C_{LFA}$, together with different strengths of repulsion, $R_{CD45TCR}$ between CD45 and TCR-pMHC. (**B**) Different repulsion strengths between CD45 and TCR-pMHC, $R_{CD45TCR}$ and CD45 and LFA-1-ICAM-1, $R_{CD45LFA}$, with $C_{LFA}$=0.06. (**C**, **D**) Amount of CD45 in the central, peripheral and distal SMAC of Figure S5A and S5B, respectively. Parameters from Table 1. Varied parameters: 18 TCR, pMHC/μm$^2$; 46 LFA-1, ICAM-1/μm$^2$. Error bars represent SD of N=10 simulations. TCR-pMHC: green, LFA-1-ICAM-1: red, CD45: white.

Movie S1. CD2-CD28 corolla in the dSMAC for $R_{CD2CD28}$=0.35μm, showing 30 minutes of IS formation.

Movie S2. Corolla pattern formation dynamics for $R_{SelfAtt}$=0.28μm, showing 30 minutes of IS formation.

Movie S3. Tracks of twenty (20) randomly selected CD2-CD58 complexes during corolla pattern formation $R_{SelfAtt}$=0.28μm, showing 30 minutes of IS formation.

Movie S4. Corolla pattern formation due to repulsive interaction between CD45 and CD2-CD58 complexes, for $R_{CD45CD2}$=0.28μm, showing 30 minutes of IS formation.
